# Estimating MIC distributions and cutoffs through mixture models: an application to establish *M. Tuberculosis* resistance

**DOI:** 10.1101/643429

**Authors:** Clara Grazian

## Abstract

Antimicrobial resistance is becoming a major threat to public health throughout the world. Researchers from around the world are attempting to contrast it by developing both new antibiotics and patient-specific treatments. It is, therefore, necessary to study these treatments, via phenotypic tests, and it is essential to have robust methods available to analyze the resistance patterns to medication, which could be applied to both new treatments and to new phenotypic tests. A general method is here proposed to study minimal inhibitory concentration (MIC) distributions and fixed breakpoints in order to separate sensible from resistant strains. The method has been applied to a new 96-well microtiter plate.

---

Public health authorities throughout the world are becoming more and more concerned about antimicrobial resistance (AMR), due to the reduced ability of standard compounds to treat infectious diseases WHO (2017); European Commission (2017); Gelband et al. (2015). Antimicrobial resistance mechanisms have been observed in bacteria Tenover (2006); Zignol et al. (2006); Kohanski et al. (2010), in fungi Perea et al. (2001); Vandeputte et al. (2011); Gulshan and Moye-Rowley (2007) and in viruses Unemo and Nicholas (2012); Yim et al. (2006); Harrigan et al. (2005).

Methods used to tackle the rise in AMR include a wiser prescription of antimicrobials, in order to develop patient-specific treatments that take into account known resistance patterns. Such patterns are studied through antimicrobial susceptibility testing (AST) to identify at which concentration of a particular drug the growth of the pathogen is inhibited. In this respect, microtiter plates allow the effectiveness of several drugs to be tested at the same time on a single clinical isolate.

Traditionally, AST methods generally rely on the definition of critical concentrations (CC), i.e. values used to differentiate between resistant and sensitive isolates which are specific to the antimicrobial agent and to the test method.

Antimicrobial data, obtained through dilution methods Wiegand et al. (2008) are registered as minimum inhibitory concentration (MIC) values, expressed in milligrams per litre (mg/l). MIC is defined as the minimal concentration of an antimicrobial substance that inhibits the visual growth of a pathogen after incubation. Since this type of test is more accurate than diffusion tests, MICs are considered the golden standard of susceptibility tests Turnidge and Paterson (2007). According to the experiment design adopted to obtain MIC values, the data of a specific drug is usually obtained in the form of a distribution.

In this paper, MIC values obtained from dilution experiments on a 96-well microtiter plate containing a liquid growth medium (broth) are analyzed, where the same dose of pathogen is cultured in each well, but in the presence of successively increasing antimicrobial concentrations. The MIC value is identified as the concentration of the first well which does not allow pathogen to grow. By convention, if no growth is observed in any well, the MIC is set to the lowest concentration available and, if growth is inhibited at each concentration level, the MIC is set to an agreed higher level of antimicrobial concentration that has not studied on the plate.

To the best of the author’s knowledge, neither clinical breakpoints nor epidemiological cutoffs have been defined for any microtiter plate. The aim of this work is to propose and compare statistical methods in order to define such thresholds.

Although the methods presented in this paper may be applied to any pathogen and any dilution method, the attention has been focused on *M. Tuberculosis*, given the importance of the resistance mechanisms developed by this pathogen. There is proof that the trend of the new cases of tuberculosis is decreasing Dheda et al. (2017), but the number of cases resistant to one or more drugs, in particular to first-line drugs (rifampicin, ethambutol, isoniazid and pyrazinamide) is increasing WHO (2015); Falzon et al. (2015). There are two main causes of the development of drug resistance, that is, either the prescription of suboptimal treatments or direct transmission.

The detection of resistance in microbiology laboratories is expected to become faster and less expensive in order to provide essential guidance to define more tailored and effective treatment schemes, and the identification of thresholds to discriminate resistant from sensitive isolates is essential. The critical concentrations of most of the anti-TB drugs have been recently revised and updated by the World Health Organisation WHO (2018), through an extensive study of the literature, and it has emerged that the identification of critical concentrations is not a simple task, as is usually assumed. However, the identification of critical concentrations is an essential step for any subsequent analysis; for instance, it could be important, in genome-wide association studies Hirschhorn and Daly (2005), to define phenotypic subgroups in order to identify the mutations associated with specific levels of resistance more clearly, in particular for those antimicrobials for which only a few resistant cases are observed (for example, when studying bedaquiline which is a new treatment for tuberculosis): in these cases, a GWAS study would describe a “wild-type” structure rather than genomic features associated with resistance.

Establishing a reliable phenotypic labeling of resistance during drug susceptibility tests is an essential step to develop reliable testing methods and to perform analysis on the identification of resistant mutations.

## 1 The dataset: resistance prediction by CRyPTIC

The CRyPTIC Consortium (Comprehensive Resistance Prediction for Tuberculosis: an International Consortium) has been created in order to collect and study around 20, 000 isolates of *M. Tuberculosis* and define a catalogue of mutations associated with resistance to 14 antituberculosis compounds: three first line drugs (isoniazid INH, rifampicin RIF, ethambutol EMB), other drugs already used in practice as antituberculosis compounds (amikacin AMI, kanamycin KAN, ethionamide ETH, phage-antibiotic synergy PAS, levofloxacin LEV, moxifloxacin MXF), two new compounds (delamanid DLM, bedaquiline BDQ) and two repurposed compounds (clofazimine CFZ, linezolid LZD).

As part of the project, the CRyPTIC Consortium have designed a UKMYC5 96-well microtiter plate. The plate has been tested by seven laboratories in Asia, Europe, South America and Africa by using 19 external quality assessment (EQA) strains, including the most studied strain of tuberculosis, H37Rv. Samples vere inoculated either 2 or 10 times and each plate was then incubated for 21 days. Two researchers from each laboratory were asked to independently identify the MIC values for each plate at four moments after incubation (day 7, day 10, day 14 and day 21) with three reading methods: Vizion imaging system, a mirrored box and an inverted-light microscope. A full description of the experiment and of the results in terms of reproducibility of the plate are available in Rancoita et al. (2018). Since the authors identified the highest level of reproducibility for readings at day 14 with the Vizion imaging system, we will focus only on data relative to this subset. Moreover, Rancoita et al. (2018) also shown that the compound PAS seems to not perform well on the plate, therefore it will be discarded in the following of the CRyPTIC study. For these reason, the outcome relative to PAS will be presented in this work, but considered more uncertain.

Notice that, as already stated, the isolates analysed in the study were subcultured several times, therefore there are repeated observations for the same sample, feature that will be considered in the model we will present. Table 1 shows the number of plates analysed for each drug, while Figure 1 shows the barplots of the MIC distributions for each compound: it is evident that some drugs show a classic bimodal distribution (RIF, RFB, INH, AMI, KAN and partially PAS), some others show a clear distribution for the sensitive cases (BDQ, CFZ, DLM), while others show a more unimodal distribution (EMB, LZD) or show distributions more spread around all the concentration range (ETH, LEV, MXF). After the validation experiment, the plate design has been changed, following the findings obtained by the analysis. We decided that the present work will use results obtained with the first plate design only, so that all the results are comparable.

**Figure 1:**
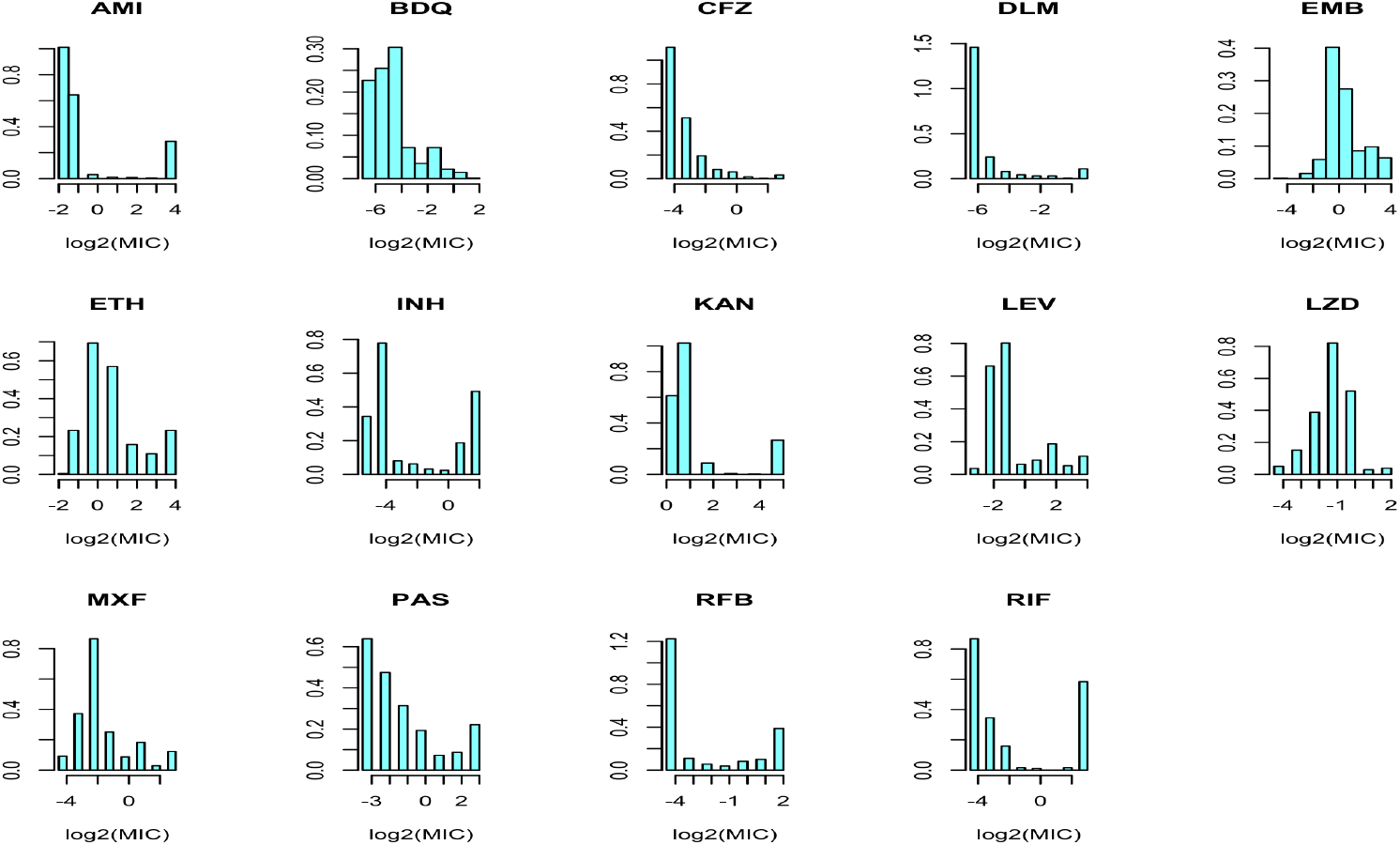
Barplots of the MIC distributions for each drug.

**Table 1:**
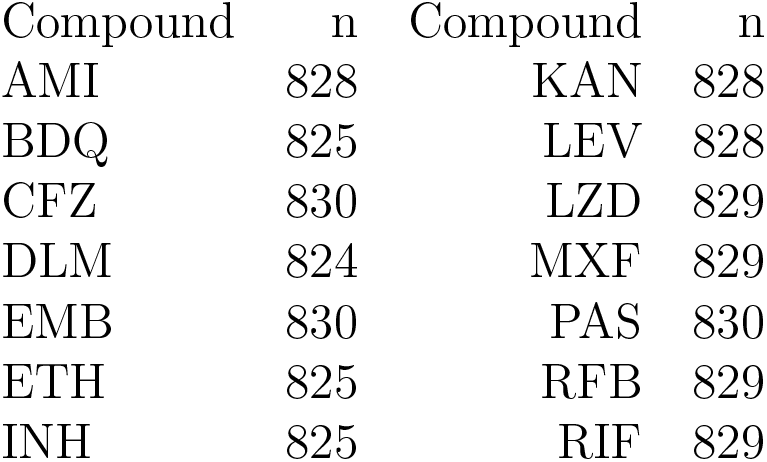
Number of isolates for each compound analysed in the CRyPTIC plate

| Compound | n | Compound | n |
| --- | --- | --- | --- |
| AMI | 828 | KAN | 828 |
| BDQ | 825 | LEV | 828 |
| CFZ | 830 | LZD | 829 |
| DLM | 824 | MXF | 829 |
| EMB | 830 | PAS | 830 |
| ETH | 825 | RFB | 829 |
| INH | 825 | RIF | 829 |

Figure 1 also shows an important feature of the dataset and, in general, of the problem of studying MIC distributions: the data are censored. Firstly, the MIC value is only partially known at the boundary of the concentration range analysed; this is particularly evident for AMI, CFZ, DLM, INH, KAN, RFB and RIF. Secondly, the MIC values are not continuous variables, they are observed at fixed level of concentrations (interval-censoring).

## 2 Statistical approaches to MIC distributions

For a single drug, defining the epidemiological cutoffs (ECOFFs) is essentially a binary classification problem, with the aim to separate the wild type isolates from the non-wild type ones, where the wild-type group includes the isolates with no acquired resistance. Visual inspection is the most straightforward way to analyze MIC data (MacGowan and Wise, 2001). From a statistical point of view, it is possible to assume that the wild-type group of isolates follows a log-normal distribution (Turnidge et al., 2006) and to identify the cutoffs by fitting a log-normal cumulative distribution by sequentially adding subsets of isolates. The optimal fit of the wild-type subpopulation is obtained when the minimal absolute difference between the estimated number of wild-type isolates and the observed number of isolates is achieved; the cutoffs is a reasonable upper limit of the wild-type distribution. More sophisticated statistical methods based on the analysis of wild-type isolates only are also available (Jaspers et al., 2014).

The empirical distributions described in Figure 1 show that the MIC values have a complex structure and standard models does not seem to appropriately fit the data. Different from the “local” methods, the “global” methods aim at modelling the whole mixing distribution (Jaspers et al., 2016): the identification of epidemiological cutoffs become a model-based classification problem. We propose to use an approach based on mixture models. These are very flexible tools used in various scientific areas, which can be used to tackle either clustering or density estimation problems. The reasons to fit a Bayesian mixture model is motivated by two aspects. First, it seems more appropriate to model the whole mixing structure rather than only the wild-type group of isolates, since the classification is unsupervised and the microtiter plates under study are characterized by biological noisiness; moreover, there exist overlapping areas in the distributions which could not be appropriately classified. The standard way to define a wild-type group is by looking at those isolates with no known conferring-resistance mutations. However, the complete patterns of resistance are not known for most of the drugs under analysis and for the new drugs are almost completely unknown. Secondly, the Bayesian framework allows for the natural incorporation of variability, taking into account biological errors in the definition of the MIC values.

Consider a random variable *Y* = (*Y*_1_,…, *Y_n_*) of size *n*. A mixture model assumes that the distribution of *Y_i_* can be written as a composition of distributions known in closed form

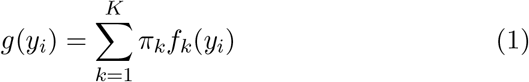

where *f_k_*(·)’s are the component probability distributions of the mixture and *π_k_*’s are known as mixture weights and are such that 0 ≤ *π_k_* ≤ 1 and 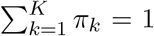. Even if the probability distributions *f_k_*(·) are free to come from any family (and they can also model either discrete or continuous random variables), in many applications it is usually assumed that all the distributions in the mixture come from the same family denoted by different parameters.

The number of component *K* is, in general, unknown and can be considered both unknown but finite (finite mixture models) or infinite (nonparametric mixtures). We refer the reader to Titterington et al. (1985) and Frühwirth-Schnatter (2006) for comprehensive books on finite mixture models and to Hjort et al. (2010) for a description of Bayesian nonparametric methods, including infinite mixture models.

This type of modelling fits well when it is assumed that there are underlying subgroups in the population and which are present with proportions *π*_1_,…, *π_K_*: in the present work, the bacterial population under analysis can be considered heterogeneous, more specifically two main subgroups can be identified, one representing the wild-type susceptible population, i.e. organisms with no acquired resistance, and one representing the non-wild-type population containing isolates which have developed some level of resistance to certain antimicrobials. However, the resistance mechanism is complicated: while the wild-type group tend to be clearly unimodal (Finch et al., 2010), and, often, considered log-normally distributed (Turnidge et al., 2006), the non-wild-type subgroup can be defined by several resistance mechanisms, which may confer different degrees of resistance. An example (Huyen et al., 2013) is the difference between mutations involving *katG* gene (associated to high level of resistance to INH - resistant to ≥ 1.6 *μ*g/ml in the UKMYC5 CRyPTIC plate) with respect to mutations involving *inhA* gene (associate to intermediate levels of resistance to INH resistant to 0.2 *μ*g/ml in the UKMYC5 CRyPTIC plate). For these reasons, we consider the number of components *K* as unknown.

The model-based classification separates the isolates in the wild-type subgroup when

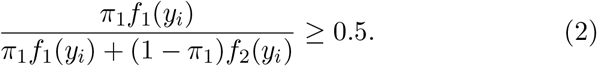

Here, the estimation of the MIC distributions is instrumental to establish cuttofs to classify the isolates into levels of resistance to the antimicrobials. It has to be briefly noted that mixture models can also be used in the general setting of density estimation, where the underlying separation in subgroups has not necessarily a biological meaning; for example, if we consider susceptibility/resistance as a gradual and continuous scale with no clear distinction between two groups, mixture models can still be used to estimate the MIC distribution.

There exist several proposals in the literature to model the MIC distributions through mixture models. The most famous is using mixture of two Gaussian distributions for the logarithm of the MIC values (Craig, 2000; Annis and Craig, 2005). However, although the MIC values can be considered ideally continuous, they are registered as discrete values, more specifically as counts for every dilution. Moreover, as already stated, considering a fixed and known number of components is a strong assumption when the mechanism of resistance are not yet fully understood.

## 3 The proposed models

We decide to implement a Bayesian analysis (Robert, 2007), so to have posterior distributions (and relative credible intervals) of all the parameters involved in the model. As part of the analysis, it is necessary to define prior distributions for all the parameters in the model, describing the prior knowledge the experimenter has about them. We have decided to use the default prior proposed in Grazian et al. (2018) for the number of components since it has been shown to have a good balance between conservativeness and accuracy and standard vague priors for the rest of the parameters as in Richardson and Green (1997), in order to reduce the influence of the prior on the posterior distribution. Bayesian estimation of mixture models is a non-standard problem, therefore Monte Carlo Markov chains (MCMC) methods are needed to approximate the posterior distribution (Robert and Casella, 2013).

Define a family of random distributions 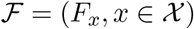, indexed by a categorical covariate *x*, which may be a vector: *x* = (*d, s*), where *d* = (1,…, 14) represents the tested drug and *s* = (1,…, 19) represents the particular strain tested.

Moreover, let **y**_1_,…, **y**_*n*_ be observations representing MIC values observed for isolates 1,…, *n*. Each isolate have been tested several times, therefore *y_xi_* represents the *i*-th repetition under condition *x*.

### Mixture of Gaussian distributions

First, we assume a Gaussian mixture model for the MIC values

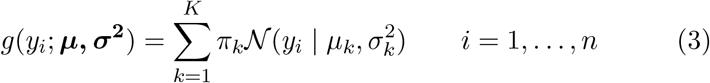

where *μ_k_* and 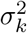 are the mean and the variance of the *k*-th component respectively and *y_i_* = *y_xi_*.

Since each strain has been subcultured several times, the MIC value related to each strain has been identified on several plates, see Table 2. Therefore, the distribution *g*(·) depends on the set of covariates *x*, specifying the tested compound and strain. In particular, we assume that

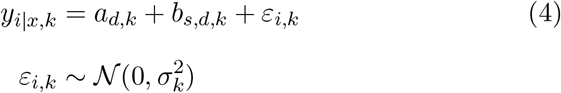

where *a_d,k_* is an intercept specific to the tested compound and *b_s,d,k_* is an intercept specific to the tested strain with respect to compound *d*.

**Table 2:**
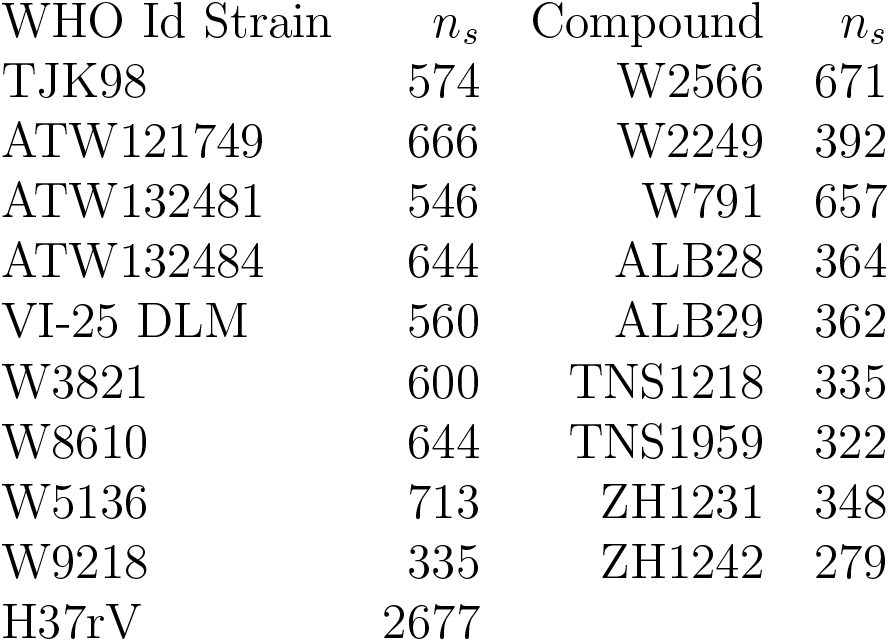
Number of plates for each isolates analysed

Equation 4 shows that a mixture model can be interpreted in a missing data framework: one could suppose the existence of a latent variable *Z_i_* taking the values in {1,…, *K*} with probabilities {*π*_1_,…, *π_K_*} and labelling the component to which the observation belongs; in other words, the conditional density of *Y_i_* diven *Z_i_* = *k* corresponds to the Gaussian distribution 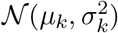. It follows that **Z**_*d*_ = (*Z*_1,*d*_,…, *Z_n_s_,d_*) is distributed according to a multinomial distribution. This latent variable representation is useful for increase the computational efficiency of MCMC algorithms (Diebolt and Robert, 1994).

This approach is extremely flexible, since it considers both the possibility that there are more than two subgroups in the data, by allowing to consider intermediate levels of resistance, and the possibility that the data are not exactly distributed as Gaussian variables: in particular, the “non-wild-type” component may be defined as a convolution of Gaussian distributions itself.

### Gaussian latent representation through mixture models

Although the previously proposed approach is very flexible, it does not take into account the censored nature of the data: MIC values are not actually continuous, they are rounded to the next two-fold dilution. Moreover, the data are essentially truncated to the minimum and to the maximum dilution chosen for the plate.

It is, then, possible to consider a mixture of distributions, where the discrete nature of the data is taken into account by rounding continuous (for instance, Gaussian) distributions. We introduce a latent variable **Y**\* ∈ ℝ which is related to the observed variable **Y** representing the registered MIC value, so that:

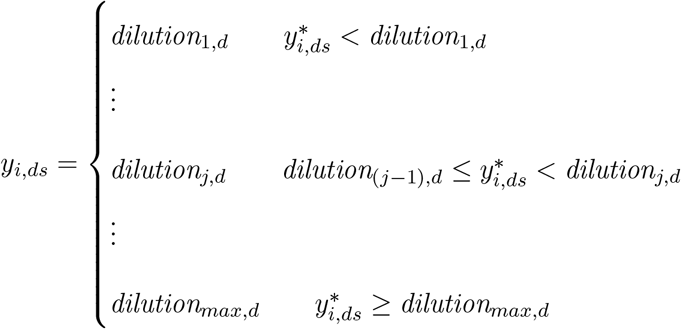

i.e. the observed *y_i_* assumes values in the set of the dilutions chosen for the drug *d* depending on a Gaussian latent variable 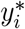 which can take value out of the set of the dilutions and which has the distribution described in Equation (3). Given a set of values representing the different dilutions 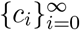 taking values in {−∞, *dilution*_1,*d*_,…, *dilution_j,d_*,…, +∞}, the probability mass function *p* for *y* is defined as

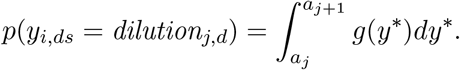

where *y*\* is assumed to follow the distribution given in (3). This approach may be seen as a latent Gaussian representation, along the line of Albert and Chib (1993). The mixing nature of the data is transferred to an implicit and richer variable which is censored and then observed only at a discrete scale.

There exist several approaches which generalize the algorithm proposed by Albert and Chib (1993) to mixture models, in particular in a nonparametric setting; for example, Kotta et al. (2005) proposes a nonparametric estimation based on infinite mixture models. Nevertheless, in this work we prefer to use a parametric mixture model with unknown number of components, in order to introduce the information that a small number of components is expected. In particular, Miller and Harrison (2014) shows an inconsistency of Dirichlet process priors to estimate the right number of components; in this respect, we have performed a separate analysis based on Dirichlet process priors, showing that the number of components has always been estimated as the maximum number of dilutions (results not shown here).

It is straightforward to define the corresponding Gibbs-sampler algorithm, which is described in Figure 2:

- *Step 1:* Generate 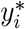 from the full conditional distributions:

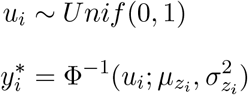
- *Step 2:* Update *K*
- *Step 3:* Update the latent variable *Z_i_* following a multinomial distribution with probabilities

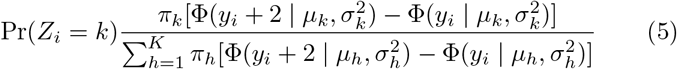
- *Step 4:*

– *Step 4a:* Update ***π***
– *Step 4a:* Update ***μ***
– *Step 4a:* Update ***σ*^2^**

**Figure 2:**
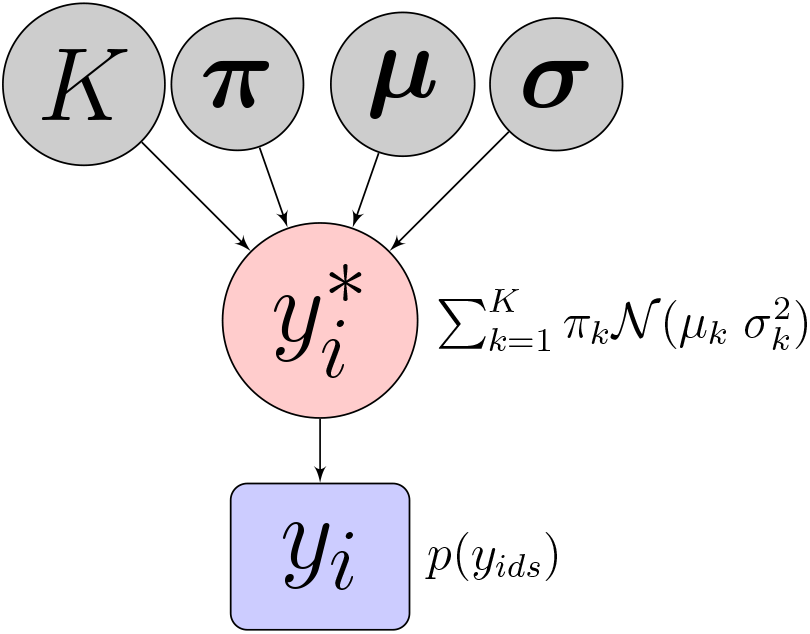
Graph representing the model: the circles are variables and the rectangle the observations. The latent variable 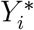 is assumed to follow a hyperprior defined as a Gaussian mixture model with parameters given in the grey circles.

## 4 Applications

The methods proposed in Section 3 is now applied to the dataset described in Section 1. The goal of the analysis is to estimate the distributions of the MIC values for each of the drugs evaluated on the microtiter plate and to identify breakpoints for the plate itself. The estimated method is based on MCMC simulations, and the evaluation of the convergence is available in the Supporting Information.

The ECOFFs identified with the censored Gaussian mixture model have been compared with three other methods: the critical concentrations identified in the recent WHO report WHO (2018) for MGIT, which is generally considered as the golden standard; the empirical cumulative distribution function approach for the wild-type group (identified as the only isolate H37Rv), as suggested by the European Committee on Antimicrobial Susceptibility Testing (EUCAST) and implemented in the ECOFFinder program EUCAST (2017), with two threshold quantiles (*q* = 0.90 and *q* = 0.95).

When a Gaussian mixture model is fitted directly to the data, the number of components is always estimated between three or four components (which are values unlikely for most of the compounds) and ECOFFs tend to be identified at lower levels (the obtained results are shown in the Supporting Information).

Figure 6 shows that the densities have been estimated correctly; an interesting property of the proposed method is that it is possible to deal with both interval censoring (related to the fact that the observations are registered at fixed dilutions) and censoring at the bounds (related to the fact that a minimal and a maximal dilution are chosen to test the isolates): the means of the components can be estimated out of the plate observational range.

Several comments can be made from an observation of Table 3. It is evident that the correct choice of the level of the quantile is essential to identify a reasonable ECOFF, and it may be changed according to the drug. However, it is *a priori* difficult to fix this level. There are drugs which clearly show a not-bimodal distribution, EMB, ETH and LZD in particular. The CRyPTIC consortium decided to change the concentrations considered in the study for these compounds, and in this way ECOFFs will be identified better in the future.

**Table 3:**
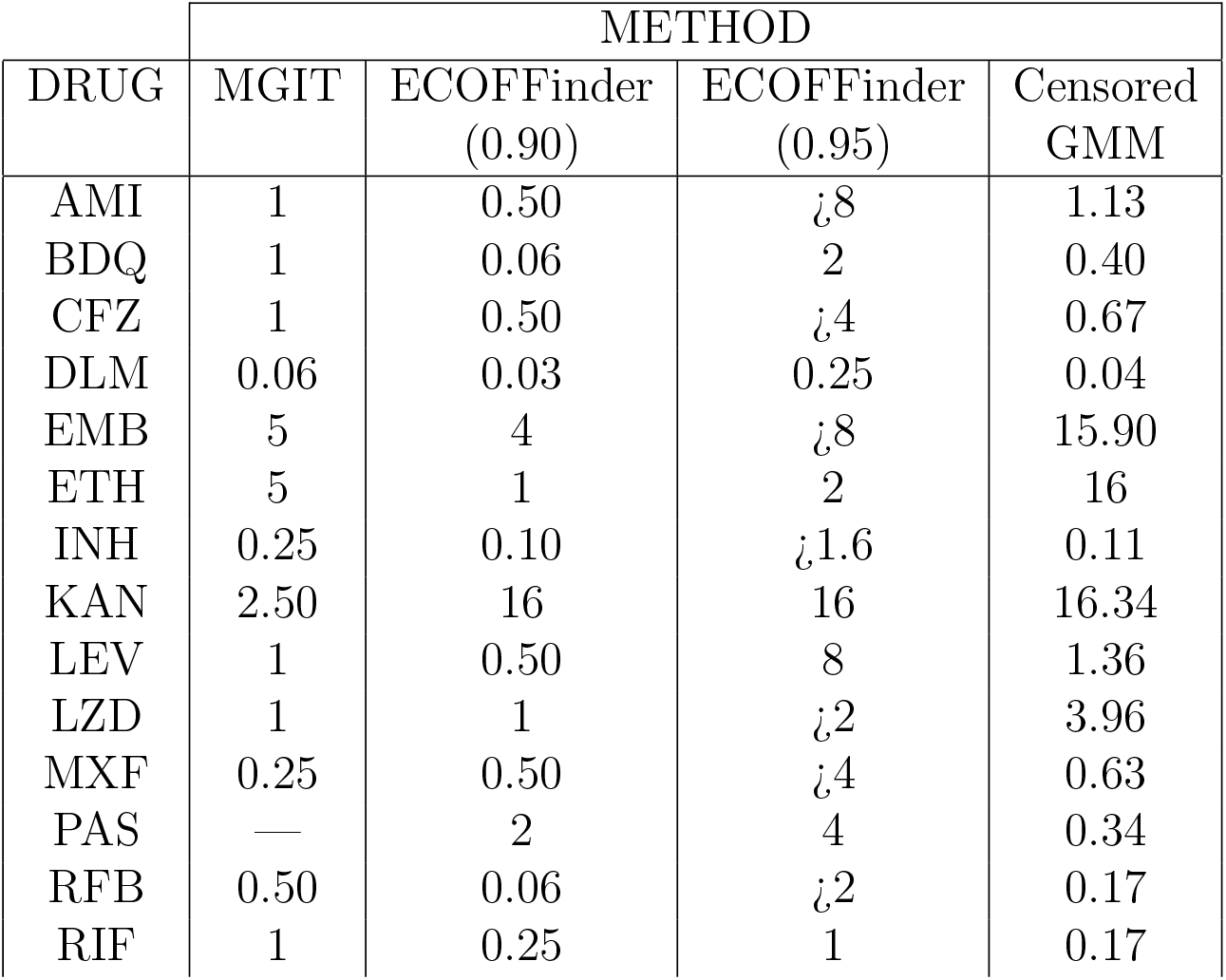
Breakpoints identified for each drug with the different methods

As already mentioned, PAS does not seem to have a recognisable distribution either, and it was removed from the study, therefore the identified ECOFF should be considered with caution.

The censored GMM and the cumulative distribution approach with a quantile of level 0.90 seems to produce similar results for the other drugs. However, the second approach is more conservative and tends to fix the critical concentrations at a lower level. The probability of identifying all the isolates that show mutations known to confer resistance to the specific drug as non-wild-type is higher for the cumulative distribution approach (Table S2 in the Supplementary Information), although it is similar to that obtained with the censored GMM for most of the cases. Nevertheless, the probability of identifying the isolates that show no mutation conferring resistance as sensitive (Table S3 in the Supplementary Information) is higher for the censored GMM. This shows that the choice of the method that should be used depends on the goal of the study: when the genomic pattern of the resistant isolates are less known, it may be better to consider a less conservative approach in order to identify those mutations that are associated more with resistance in a clear way: this is the case of BDQ, CFZ and DLM. When the goal is to analyze the unknown mutations associated with an intermediate level of resistance in drugs that have already been studied, it could be useful to apply different methods and then to compare the results.

## 5 Conclusions

A general method has been proposed to approximate MIC distributions and identify critical concentrations. This method may be applied to several testing situations and in the presence or absence of repeated tests on the same isolates.

**Figure 3:**
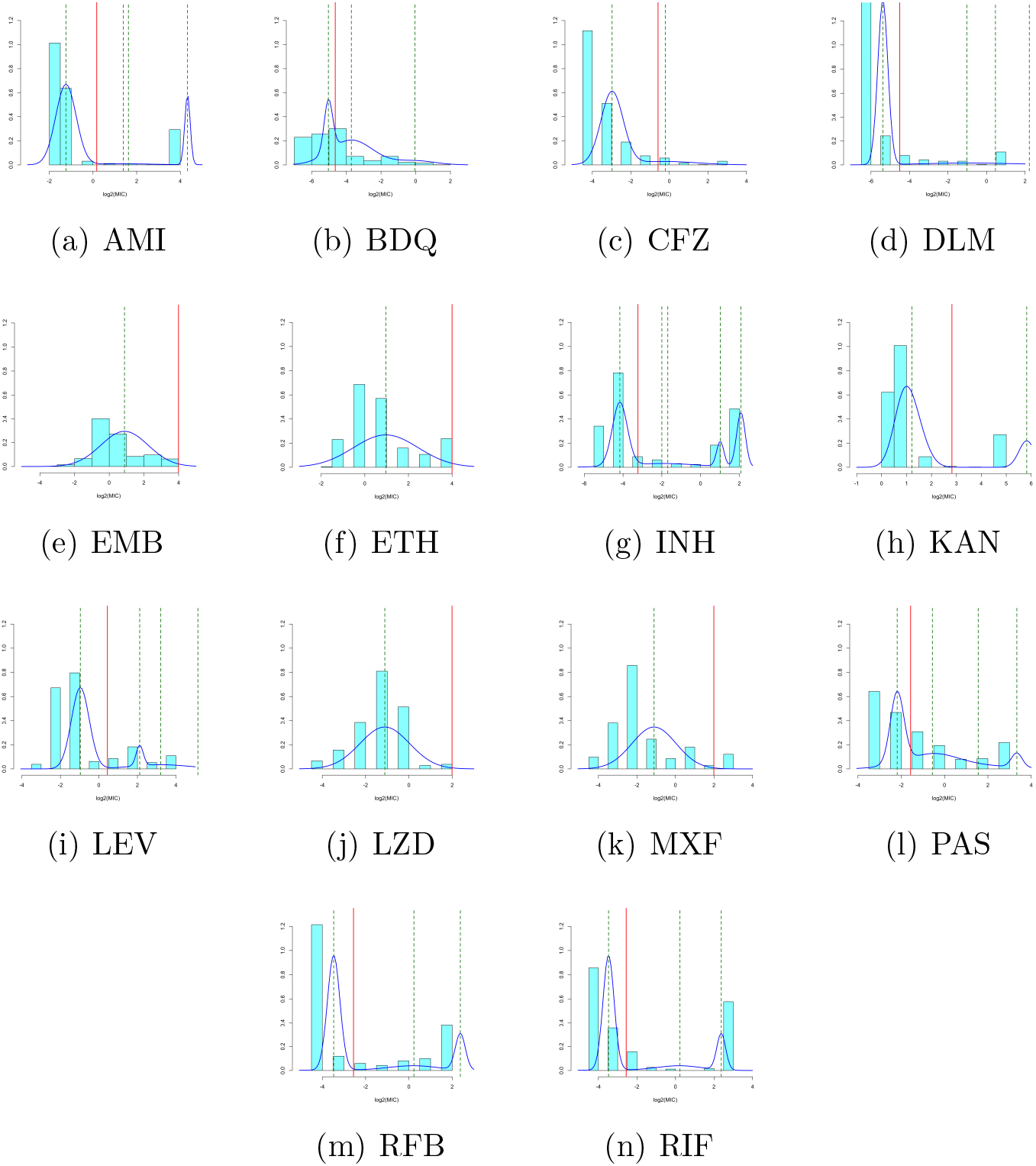
Identification of ECOFFs using a latent Gaussiam mixture model: the blue line stands for estimated density, the locations of each component are represented in green and the breakpoint is represented in red.

The proposed method has the advantage of reducing the number of necessary assumptions and inputs. Approaches based on the definition of a wild-type group, such as the cumulative distribution approach, can be influenced to a great extent by this definition, which is particularly sensitive when little is known about the genomic resistance pattern or when the antimicrobials have been available for a long period of time. This is the case of *M. Tuberculosis*: the first-line drugs were introduced in the 1950s and 1960s, while the genes related to the resistance to the new or repurposed drugs, such as bedaquiline, clofazimine, linezolid and delamanid, are still unknown.

The approach has been compared with other available approaches considering a dataset developed within the CRyPTIC project. The study has included data that were collected during the first phase of the project in order to validate the microtiter plate. The results show that the proposed approach leads to the highest combination of sensitivity and specificity for all of the considered drugs, as far as the known genomic patterns of resistance are concerned.

At the moment, the dataset is somewhat limited. However this is just the first attempt to define ECOFFs for the microtiter plate used in the CRyPTIC project. As long as more data are collected and more genomic mutations related to resistance will be identified, the calibration of the breakpoints will be improved.

## Acknoledgements

This work has partially been developed under the PRIN2015 supported-project: Environmental processes and human activities: capturing their interactions via statistical methods (EPHAStat), funded by MIUR (Italian Ministry of Education, University and Scientific Research) (20154X8K23-SH3).

## Appendix 1: Study of convergence for the mixture of Gaussian distributions model

**Figure 4:**
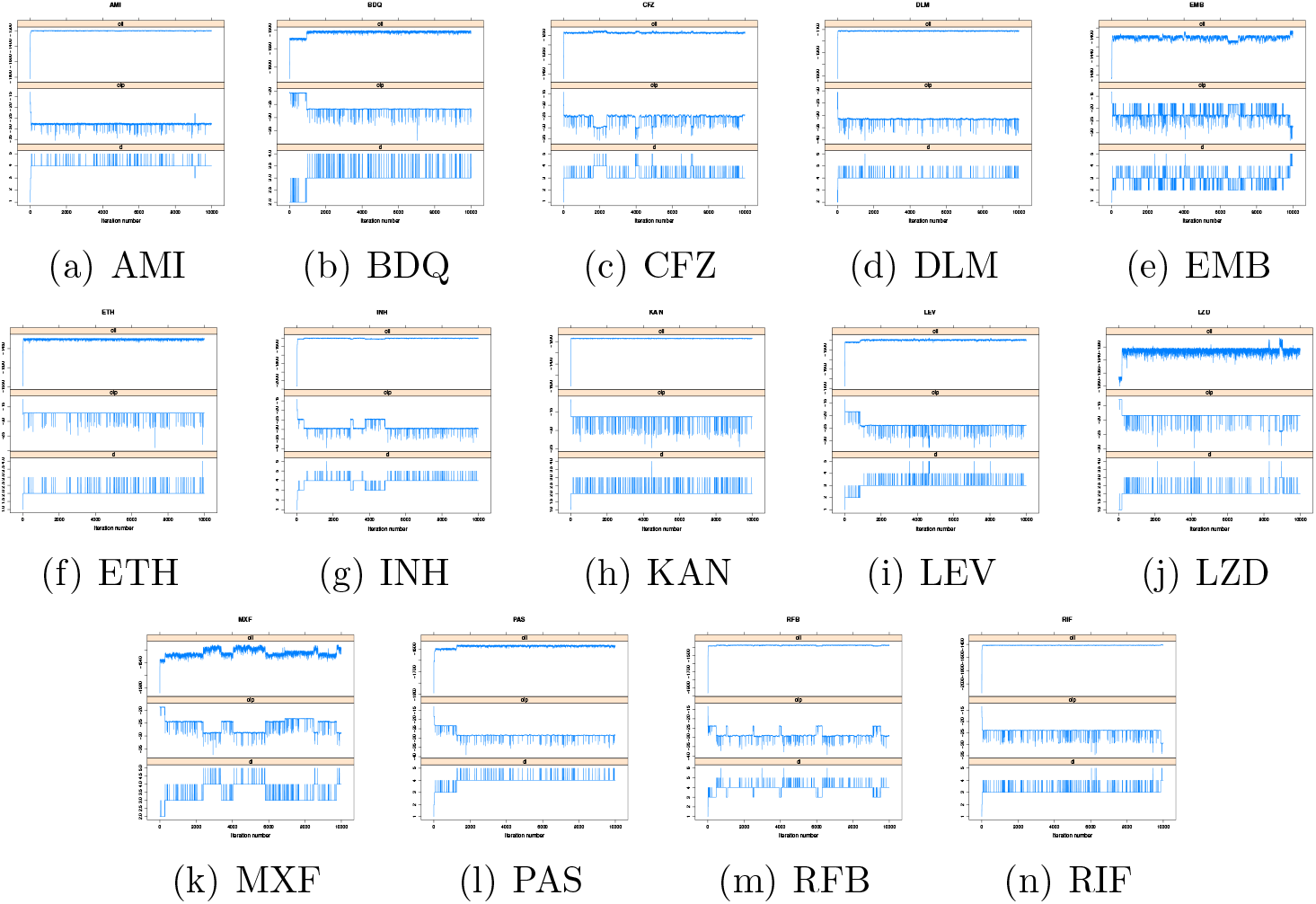
Convergence for each drug: in each figure the MCMC chains are shown relative to the likelihood function of the accepted values (above), the prior distribution of the accepted values (middle) and the chain of the number of components (below).

## Appendix 2: Study of convergence for the mixture of Gaussian distributions model

**Figure 5:**
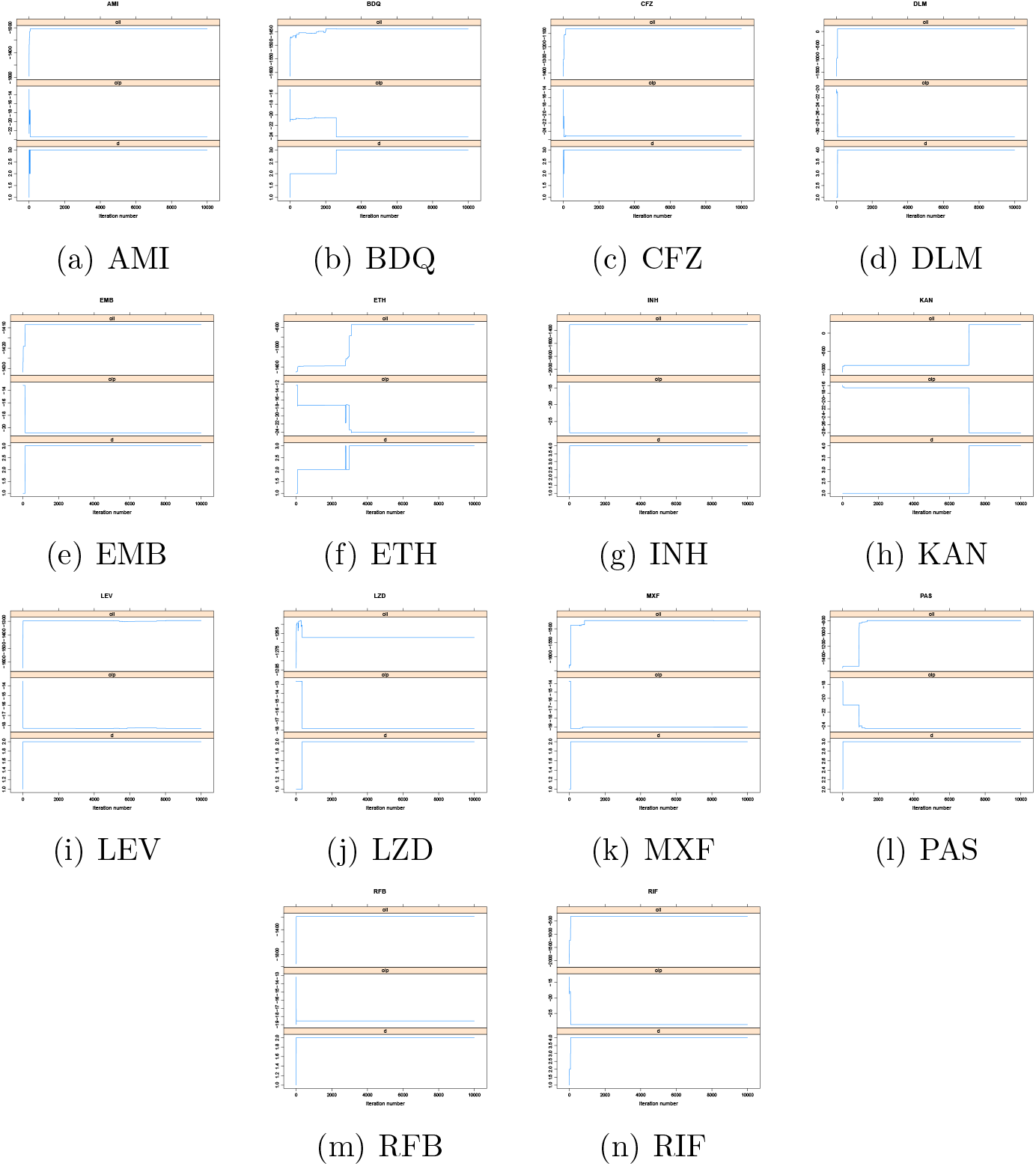
Convergence for each drug: in each figure the MCMC chains are shown relative to the likelihood function of the accepted values (above), the prior distribution of the accepted values (middle) and the chain of the number of components (below).

## Appendix 3: Definition of ECOFFs with Gaussian mixture models

**Figure 6:**
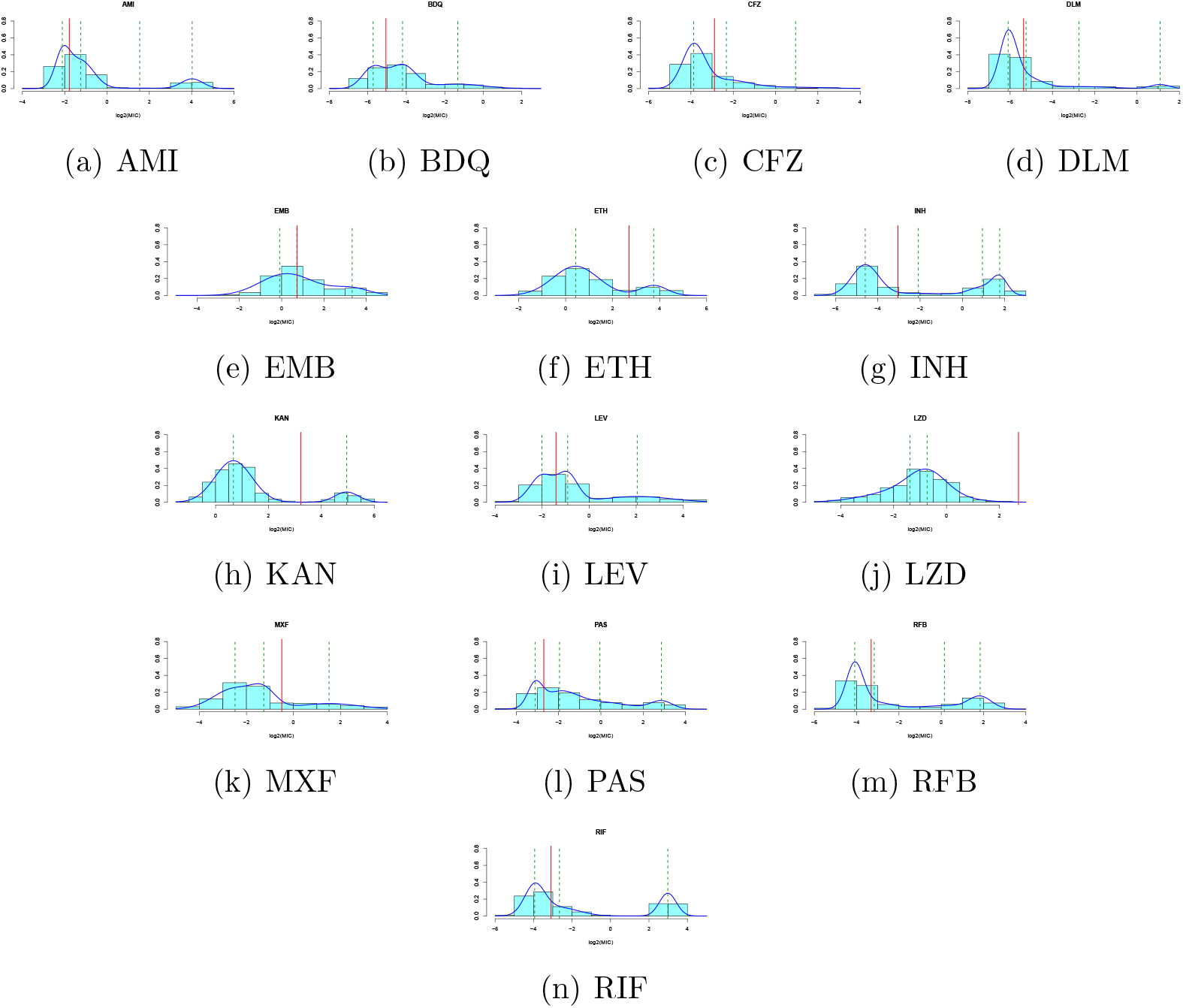
Identification of ECOFFs by using a Gaussiam mixture model: the estimated density is the blue line, the means of each components are represented in green and the breakpoint is represented in red.

## Appendix 4: Genotype of the strains

**Table 4:** Genotype of the strains under analysis for each drug.

| Strain | MGIT | Gene | Mutation | Drug |
| --- | --- | --- | --- | --- |
| H37rV | S | - | - | - |
| TJK 98 | R | ddn | Trp88* | Delamanid |
|  | R | katG | Ser315Thr | Isoniazid |
|  | R | rpoB | Ser450Leu | Rifabutin |
|  | R | rpoB | Ser450Leu | Rifampicin |
| ATW121749 | S | - | - | - |
| ATW132481 | R | Rv0678 | DelA-Gln115-fs | Bedaquiline |
|  | R | Rv0678 | DelA-Gln115-fs | Clofazimine |
| ATW132484 | R | Rv0678 | Tyr92* | Bedaquiline |
|  | R | Rv0678 | Tyr92* | Clofazimine |
| VI-25 DLM | R | rrs | a1401g | Amikacin |
|  | R | embB | Met306Val | Ethambutol |
|  | R | ethA | DelT-Lys37-fs | Ethionamide |
|  | R | prom fabG1-inhA | -34 c - Del | Ethionamide |
|  | R | katG | Ser315Thr | Isoniazid |
|  | R | prom fabG1-inhA | -34 c - Del | Isoniazid |
|  | R | rrs | a1401g | Kanamycin |
|  | R | gyrA | Ala90Val | Levofloxacin |
|  | R | gyrA | Ala90Val | Moxifloxacin |
|  | R | rpoB | Ala90Val | Rifabutin |
|  | R | rpoB | Ala90Val | Rifampicin |
| W3821 | R | ethA | Asp357Tyr | Ethionamide |
|  | R | inhA | Asn231Asp | Ethionamide |
|  | R | katG | Ser315Thr | Isoniazid |
|  | R | gyrA | Ala90Val | Levofloxacin |
|  | R | gyrA | Ser91Pro | Levofloxacin |
|  | R | gyrA | Ala90Val | Moxifloxacin |
|  | R | gyrA | Ser91Pro | Moxifloxacin |
|  | R | rpoB | Ser450Leu | Rifabutin |
|  | R | rpoB | Ser450Leu | Rifampicin |
| W8610 | R | rrs | a1401g | Amikacin |
|  | R | rrs | a1401g | Kanamycin |
| W5136 | R | katG | Ser315Thr | Isoniazid |
| W9218 | R | rrs | a1401g | Amikacin |
|  | R | embB | Met306Ile | Ethambutol |
|  | R | katG | Ser315Thr | Isoniazid |
|  | R | rrs | a1401g | Kanamycin |
|  | R | rpoB | Ser450Leu | Rifabutin |
|  | R | rpoB | Ser450Leu | Rifampicin |
| W2566 | R | embB | Met306Ile | Ethambutol |
|  | R | katG | Ser315Thr | Isoniazid |
|  | R | gyrA | Ala90Val | Levofloxacin |
|  | R | gyrA | Ala90Val | Moxifloxacin |
|  | R | rpoB | Ser450Leu | Rifabutin |
|  | R | rpoB | Ser450Leu | Rifampicin |
| W2249 | S | - | - | - |
| W791 | R | gyrA | Ala90Val | Levofloxacin |
|  | R | gyrA | Ala90Val | Moxifloxacin |
| ALB28 | S | - | - | - |
| ALB29 | S | - | - | - |
| TNS1218 | R | rpoB | Ser450Leu | Rifabutin |
|  | R | rpoB | Ser450Leu | Rifampicin |
| TNS1959 | R | katG | Ser315Thr | Isoniazid |
|  | R | rpoB | Ser450Leu | Rifabutin |
|  | R | rpoB | Ser450Leu | Rifampicin |
| ZH1231 | R | prom fabG1-inhA | -15c- > t | Ethionamide |
|  | R | prom fabG1-inhA | -15c- > t | Isoniazid |
| ZH1242 | R | katG | Ser315Thr | Isoniazid |
|  | R | thyA | Val261Gly | PAS |

## Appendix 5: Sensitivity and Specificity

**Table 5:** Percentage of isolates showing known mutations that confer resistance to a specific drug and which have been identified as resistant by a specific method.

| DRUG | METHOD |  |  |
| --- | --- | --- | --- |
|  | ECOFFinder (.90) | ECOFFinder (.95) | Censored GMM |
| AMI | 100.00 | 0.00 | 100.00 |
| BDQ | 55.19 | 0.00 | 40.44 |
| CFZ | 14.21 | 0.00 | 14.21 |
| DLM | 100.00 | 62.20 | 96.34 |
| EMB | 73.01 | 0.00 | 25.77 |
| ETH | 86.59 | 68.90 | 0.00 |
| INH | 94.10 | 0.00 | 94.10 |
| KAN | 100.00 | 100.00 | 99.12 |
| LEV | 91.40 | 20.43 | 88.71 |
| LZD | 1.35 | 0.00 | 0.00 |
| MXF | 85.95 | 0.00 | 85.95 |
| PAS | 50.85 | 44.07 | 76.27 |
| RFB | 100.00 | 0.00 | 97.63 |
| RIF | 97.63 | 96.05 | 97.63 |

**Table 6:**
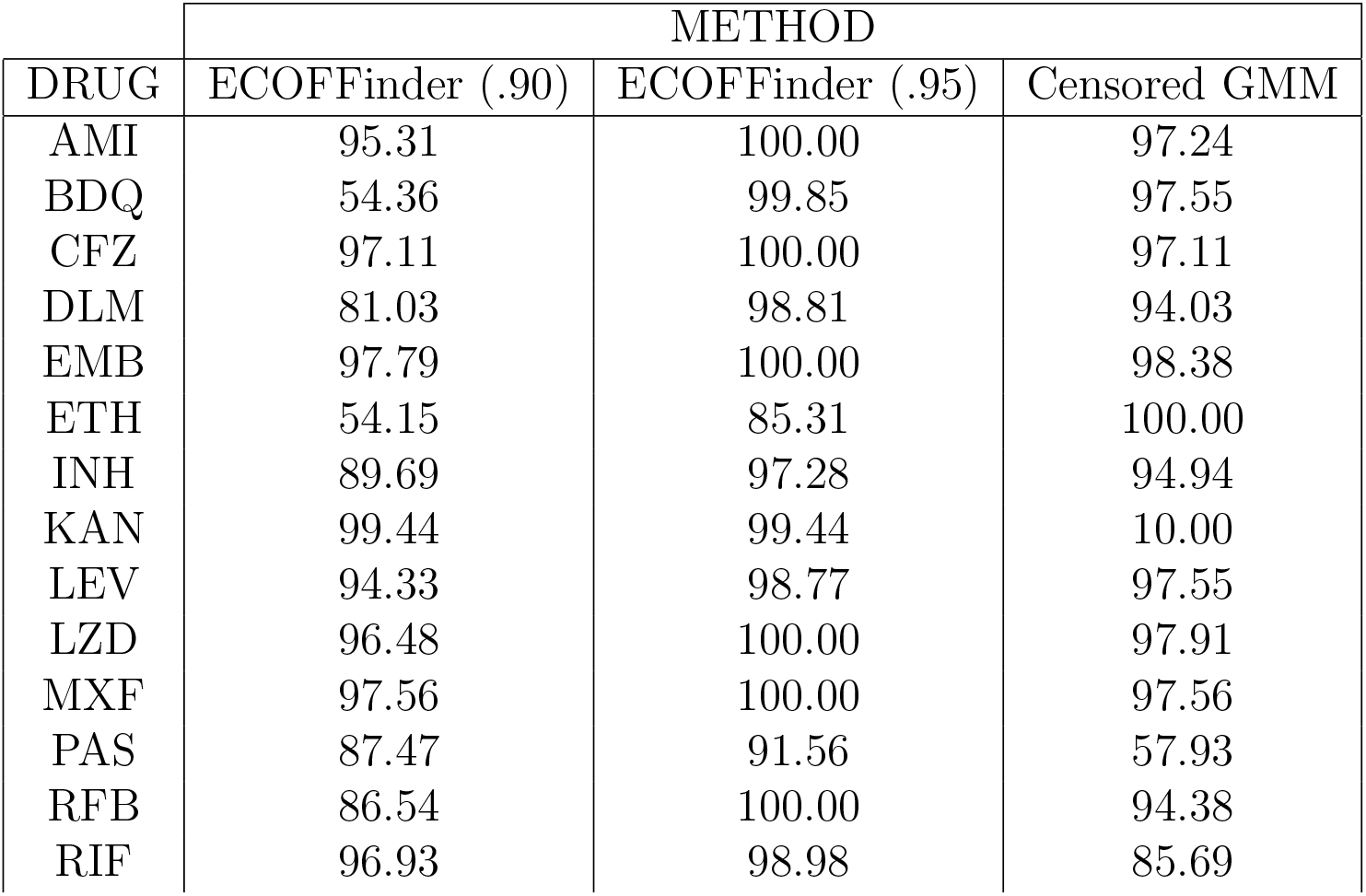
Percentage of isolates showing no known mutation that confer resistance to a specific drug and which have been identified as sensitive by a specific method.

## References

WHO (2017) Global framework for development & stewardship to combat antimicrobial resistance? World Health Organization: Geneva, Switzerland.

European Commission (2017) A European one health action plan against antimicrobial resistance (AMR). European Commission.

Gelband H, et al. (2015) The state of the world’s antibiotics 2015. Wound Healing Southern Africa 8(2):30–34.

Tenover FC (2006) Mechanisms of antimicrobial resistance in bacteria. American journal of infection control 34(5):S3–S10.

Zignol M, et al. (2006) Global incidence of multidrug-resistant tuberculosis. The Journal of infectious diseases 194(4):479–485.

Kohanski MA, DePristo MA, Collins JJ (2010) Sublethal antibiotic treatment leads to multidrug resistance via radical-induced mutagenesis. Molecular cell 37(3):311–320.

Perea S, et al. (2001) Prevalence of molecular mechanisms of resistance to azole antifungal agents in candida albicans strains displaying high-level fluconazole resistance isolated from human immunodeficiency virus-infected patients.

Vandeputte P, Ferrari S, Coste AT (2011) Antifungal resistance and new strategies to control fungal infections. International journal of microbiology 2012.

Gulshan K, Moye-Rowley WS (2007) Multidrug resistance in fungi. Eukaryotic Cell 6(11):1933–1942.

Unemo M, Nicholas RA (2012) Emergence of multidrug-resistant, extensively drug-resistant and untreatable gonorrhea. Future microbiology 7(12):1401–1422.

Yim HJ, et al. (2006) Evolution of multi-drug resistant hepatitis B virus during sequential therapy. Hepatology 44(3):703–712.

Harrigan PR, et al. (2005) Predictors of hiv drug-resistance mutations in a large antiretroviral-naive cohort initiating triple antiretroviral therapy. The Journal of infectious diseases 191(3):339–347.

Wiegand I, Hilpert K, Hancock RE (2008) Agar and broth dilution methods to determine the minimal inhibitory concentration (MIC) of antimicrobial substances. Nature protocols 3(2):163.

Turnidge J, Paterson DL (2007) Setting and revising antibacterial susceptibility breakpoints. Clinical microbiology reviews 20(3):391–408.

Dheda K, et al. (2017) The epidemiology, pathogenesis, transmission, diagnosis, and management of multidrug-resistant, extensively drug-resistant, and incurable tuberculosis. The Lancet Respiratory medicine 5(4):291–360.

WHO (2015) Global tuberculosis report 2015. (World Health Organization).

Falzon D, et al. (2015) Multidrug-resistant tuberculosis around the world: what progress has been made? European Respiratory Journal 45(1):150–160.

WHO (2018) Technical report on critical concentrations for drug susceptibility testing of medicines used in the treatment of drug-resistant tuberculosis, (World Health Organization), Technical report.

Hirschhorn JN, Daly MJ (2005) Genome-wide association studies for common diseases and complex traits. Nature Reviews Genetics 6(2):95.

Rancoita PM, et al. (2018) Validating a 14-drug microtitre plate containing bedaquiline and delamanid for large-scale research susceptibility testing of mycobacterium tuberculosis. bioRxiv p. 244731.

MacGowan AP, Wise R (2001) Establishing mic breakpoints and the interpretation of in vitro susceptibility tests. Journal of Antimicrobial Chemotherapy 48(suppl_1):17–28.

Turnidge J, Kahlmeter G, Kronvall G (2006) Statistical characterisation of bacterial wild-type mic value distributions and the determination of epidemiological cut-off values. Clinical Microbiology and Infection 12(5):418–425.

Jaspers S, Aerts M, Verbeke G, Beloeil PA (2014) Estimation of the wild-type minimum inhibitory concentration value distribution. Statistics in medicine 33(2):289–303.

Jaspers S, Lambert P, Aerts M,, et al. (2016) A Bayesian approach to the semiparametric estimation of a minimum inhibitory concentration distribution. The Annals of Applied Statistics 10(2):906–924.

Titterington DM, Smith AF, Makov UE (1985) Statistical analysis of finite mixture distributions. (Wiley,).

Frühwirth-Schnatter S (2006) Finite mixture and Markov switching models. (Springer Science & Business Media).

Hjort NL, Holmes C, Müller P, Walker SG (2010) Bayesian nonparametrics. (Cambridge University Press) Vol. 28.

Finch RG, Greenwood D, Whitley RJ, Norrby SR (2010) Antibiotic and chemotherapy e-book. (Elsevier Health Sciences).

Huyen MN, et al. (2013) Isoniazid resistance mutations: epidemiology and effect on tuberculosis treatment outcomes. Antimicrobial agents and chemotherapy pp. AAC–00077.

Craig BA (2000) Modeling approach to diameter breakpoint determination. Diagnostic microbiology and infectious disease 36(3):193–202.

Annis DH, Craig BA (2005) Statistical properties and inference of the antimicrobial mic test. Statistics in medicine 24(23):3631–3644.

Robert C (2007) The Bayesian choice: from decision-theoretic foundations to computational implementation. (Springer Science & Business Media).

Grazian C, Villa C, Liseo B (2018) On a loss-based prior for the number of components in mixture models. arXiv preprint arXiv:1807.07874.

Richardson S, Green PJ (1997) On Bayesian analysis of mixtures with an unknown number of components (with discussion). Journal of the Royal Statistical Society: series B (statistical methodology) 59(4):731–792.

Robert C, Casella G (2013) Monte Carlo statistical methods. (Springer Science & Business Media).

Diebolt J, Robert CP (1994) Estimation of finite mixture distributions through Bayesian sampling. Journal of the Royal Statistical Society. Series B (Methodological) pp. 363–375.

Albert JH, Chib S (1993) Bayesian analysis of binary and polychotomous response data. Journal of the American statistical Association 88(422):669–679.

Kottas A, Müller P, Quintana F (2005) Nonparametric Bayesian modeling for multivariate ordinal data. Journal of Computational and Graphical Statistics 14(3):610–625.

Miller JW, Harrison MT (2014) Inconsistency of Pitman-Yor process mixtures for the number of components. The Journal of Machine Learning Research 15(1):3333–3370.

EUCAST (2017) Eucast sop 1.2: Setting breakpoints for new agents, (EUCAST), Technical report.

